# A novel *f*-divergence based generative adversarial imputation method for scRNA-seq data analysis

**DOI:** 10.1101/2023.08.28.555223

**Authors:** Tong Si, Zackary Hopkins, John Yanev, Jie Hou, Haijun Gong

## Abstract

Comprehensive analysis of single-cell RNA sequencing (scRNA-seq) data can enhance our understanding of cellular diversity and aid in the development of personalized therapies for individuals. The abundance of missing values, known as dropouts, makes the analysis of scRNA-seq data a challenging task. Most traditional methods made assumptions about specific distributions for missing values, which limit their capability to capture the intricacy of high-dimensional scRNA-seq data. Moreover, the imputation performance of traditional methods decreases with higher missing rates. We propose a novel *f* -divergence based generative adversarial imputation method, called sc-*f* GAIN, for the scRNA-seq data imputation. Our studies identify four *f* -divergence functions, namely cross-entropy, Kullback-Leibler (KL), reverse KL, and Jensen-Shannon, that can be effectively integrated with the generative adversarial imputation network to generate imputed values without any assumptions, and mathematically prove that the distribution of imputed data using sc-*f* GAIN algorithm is same as the distribution of original data. Real scRNA-seq data analysis has shown that, compared to many traditional methods, the imputed values generated by sc-*f* GAIN algorithm have a smaller root-mean-square error, and it is robust to varying missing rates, moreover, it can reduce imputation bias. The flexibility offered by the *f* -divergence allows the sc-*f* GAIN method to accommodate various types of data, making it a more universal approach for imputing missing values of scRNA-seq data.

## Introduction

The genomics and transcriptomics studies have been revolutionized by the swift advancements in single-cell RNA sequencing (scRNA-seq) technology, which enable researchers to simultaneously profile the transcriptomes of thousands of individual cells [1, 2], providing a comprehensive view of the cellular heterogeneity within a tissue or organism [3]. Most of studies [4, 5, 6, 7, 8, 9] are based on the traditional bulk RNA-seq or microarray experiments, which calculate the mean gene expression profile of cells in a sample, ignoring the heterogeneity and genomic variability among individual cells. Clinical studies have found that many drugs are not effective to some patients due to the cellular diversity or heterogeneous effects across individuals [10]. The scRNA-seq data can help identify distinct cell types that may have different functions or respond differently to the same stimuli or treatment [11]. scRNA-seq data can also be used to reconstruct cell-type-specific regulatory networks [12] and identify important regulatory processes or patterns that correspond to specific cell types or processes. Comprehensive analysis of scRNA-seq data can improve our understanding of cellular heterogeneity and disease mechanisms, and potentially help develop personalized and targeted treatments for individual patients based on their unique cellular profiles [13].

The missing values, which accounts for more than 50% and sometimes over 90% of the scRNA-seq data [14, 15], are often represented as 0 or values close to 0, making the scRNA-seq data analysis a challenging task [16]. Prevalence of missing values causes inaccurate reconstruction of cell-type-specific networks, consequently limits the potential power and benefits of single-cell RNA sequencing technologies [17, 18]. For instance, the missing data could get confounded with some genuinely low captured data, leading to a failure in identifying the regulatory functions [16] and deciphering the underlying biological mechanisms of specific cells.

Numerous computational techniques [19, 20, 21, 22, 23, 24, 25] have been developed to impute the missing values in the scRNA-seq data. Although some methods have shown efficacy in recovering missing values in small-scale scRNA-seq data, each method faces different challenges [26]. Markov Affinity-based Graph Imputation of Cells (MAGIC) [21] method creates a Markov transition matrix using the similarity matrix of single cells described by the Pearson correlation or mutual information to impute missing values. However, the MAGIC method necessitates the use of information from analogous cells or the consolidation of genes derived from scRNA-seq data that has been observed. This can result in inaccurate estimation of cell variability and reduced gene ranking performance [26], as well as potential oversmoothing of the data. SAVER [22] introduces different prior distributions to model the observed data, then develops a Bayesian-based model to impute missing values for each cell. Due to its dependence on a Markov Chain Monte Carlo algorithm for adjusting all parameters, its computational expenses are significant and may render it unscalable for extensive datasets [27]. PBLR [23] is a bounded low-rank recovery imputation method that constructs a consensus matrix and employs hierarchical clustering to identify cell sub-populations and submatrices for imputation. Another popular method, scImpute [14] assumes majority of the genetic information can be captured by a smaller number of latent factors, and utilizes an autoencoder to approximate the absent values. Most of these traditional methods [23, 22, 20, 28, 29] assume the missing values follow some specific distributions, so they could not accurately capture the intricacy of the high-dimensional scRNA-seq data. Furthermore, the imputation accuracy of these methods will decrease as the missing rate increases.

The outstanding performance of deep learning methods have attracted lot of research attention in the field of computational biology [24]. Deep generative methods, such as generative adversarial networks (GANs) [30], have emerged as powerful techniques for learning models from real-world data [31, 32]. GANs have demonstrated exceptional proficiency in diverse domains such as generating images [30] and videos [33], as well as image inpainting, where the missing parts of a corrupted image are filled in based on the remaining portion with outstanding performance. Recently, a new imputation method called GAIN, also known as generative adversarial imputation nets [34], was proposed to recover missing values. GAIN uses a vanilla GAN architecture with a generator network to impute missing values while a discriminator network evaluates the quality of the imputed values. GAIN has shown promising results in imputing missing values with an adversarial loss described by a binary cross entropy (BCE) function. However, there are still some limitations associated with this method. For instance, the use of the cross-entropy loss function may be inadequate for some intricate distributions. Moreover, the original GAN architecture that GAIN used has a serious problem called mode collapse. One possible solution to address these problems is to incorporate *f* -divergence in the GAN’s architecture.

In this work, we are motivated to develop a novel imputation approach that does not rely on distribution assumptions, called sc-*f* GAIN, standing for *f* -divergence based generative adversarial imputation network, for the scRNA-seq data analysis. Our work has two major novelties compared with vanilla GAIN and other existing methods. For the first time, our studies identify four *f* -divergence functions, namely cross-entropy, Kullback-Leibler (KL), reverse KL, and Jensen-Shannon, that can be integrated with GAIN to generate imputed values without any assumptions, and mathematically prove that the distribution of imputed data using sc-*f* GAIN algorithm is same as the distribution of original data. Finally, we evaluated the effectiveness and potential limitations of sc-*f* GAIN method by training it on real scRNA-seq data and comparing it to other imputation methods.

## Materials and Methods

Generative modeling is a technique to learn the underlying probability distribution of a high-dimensional real-world dataset. Generative adversarial networks (GANs) are a class of deep generative models which use two competing neural networks to produce new data that resembles the real-world training data which follows an unknown probability distribution (*x∼p*(*x*)) [30]. These two networks are called Generator (*G*(*θ*)) and Discriminator (*D*(*ϕ*)), which are implemented as deep neural networks. The objective of the Generator is to transform a noise vector (*z ∼q*(*z*)) to generate synthetic/fabricated samples (*G*(*z, θ*)) that are indistinguishable from authentic samples to trick the Discriminator. On the contrary, the role of the Discriminator is to differentiate the authentic samples (*x*) from the fabricated ones (*G*(*z, θ*)) produced by the Generator. *θ* and *ϕ* are the parameters of the Generator and Discriminator networks, respectively, and they are learned using Backpropagation algorithm by minimizing the cross entropy loss functions in the vanilla GANs [30]. The primary objective of the vanilla GAN is to minimize the generator loss, meanwhile maximize the discriminator loss, which is equivalent to solving the minimax optimization problem:

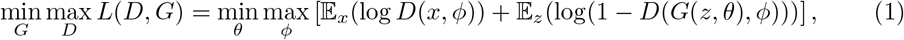

Model collapse is a serious problem in the vanilla GANs where the generator network produces limited variations of samples that do not capture the diversity of the true data distribution, resulting in the discriminator network becoming too skilled at discriminating them [35]. One reason for mode collapse in vanilla GANs is the binary cross-entropy loss, which is not suitable for some complex distributions and might not capture the high-dimensional structure of the data. One possible solution to address this problem is to replace cross-entropy loss by the *f* -divergence based loss function to mitigate the mode collapse problem in GANs. *f* -divergence is a family of statistical measures which quantify the discrepancy between two distributions, with a higher value indicating greater dissimilarity between the generated samples and the actual data. Next we will briefly introduce the *f* -GAN [36] method.

### *f* -GAN: *f* -divergence based Generative Adversarial Networks

*f* -GAN, or the *f* -divergence based generative adversarial network, is a variant of generative adversarial networks with the loss function change to the class of *f* -divergence, which is a generalization of Kullback-Leibler (KL) divergence and includes other divergences such as reverse KL, Jensen-Shannon as special cases. The *f* -GAN framework was first proposed in Nowozin *et al*.’s work [36] to generate synthetic pictures. To provide a better understanding of this framework, we will briefly review some fundamental concepts related to *f* -divergence and *f* -GAN.

The *f* -divergence, also known as the Ali-Silvey distances [37], between two probability distributions *P* and *Q* is defined as:

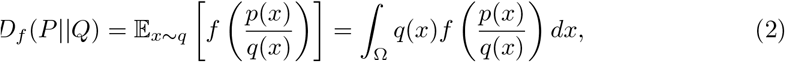

where, *p*(*x*) and *q*(*x*) are two continuous probability density function on the domain Ω, and the generator function *f* is a proper, lower-semicontinuous, convex generator function with *f* (1) = 0. Since the *f* -divergence measures the difference between two distributions, so, it could be used as a measure of the loss functions in the GANs.

The estimation of *f* -divergence need use the convex conjugate of a convex function. Every convex lower-semicontinuous function *f* has a convex conjugate function *f*\*, which is also known as Fenchel-Legendre transform of *f*. The convex conjugate is defined to be [38]: 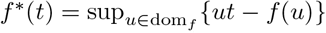. It has been proved that *f*\* is also convext and lower-semicontinuous if *f* (*x*) is a convex function on ℝ. The lower bound of the *f* -divergence can be estimated using the Fenchel conjugate function, which is expressed as [36]:

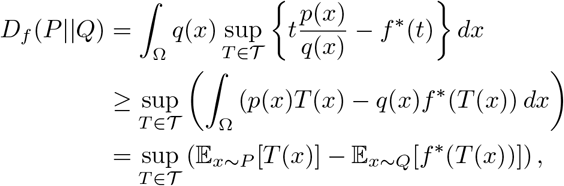

where, *T* (*x*) is any Borel function. After taking the variation with respect to *T*, we get 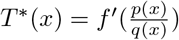 which can be used to choose the generator function *f* and design the class of functions *T*.

Given a convex function *f*, the objective of *f* -GAN is to minimize the *f* –divergence *D*_*f*_ (*P*‖*Q*), that is, solve the following minimax problem using the variational divergence minimization (VDM) method:

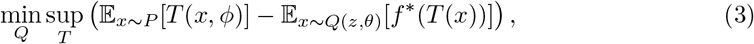

where, *Q*(*z, θ*) is a generator implemented by a neural network, which takes a random vector *z* as input and output a sample of interest. The Borel function *T* (*x*) is called a critic function, which is also approximated by a neural network. If *S*_*ϕ*_ is the output of neural network with parameters *ϕ, g*_*f*_ is an output activation function specific to the choice of *f* -divergence, then we can have the critic function being clarified as *T* (*x*) = *g*_*f*_ (*S*_*ϕ*_ (*x*)), and the above minimax problem can be expressed as

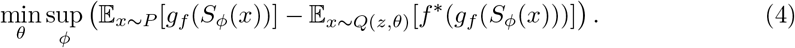

By using a variety of *f* functions and output activation functions *g*_*f*_, we can obtain various *f* -divergence [36], including the divergence of cross entropy (CE), Kullback-Leibler (FKL), Reverse KL (RKL), Jensen-Shannon (JS), Pearson *χ*^2^ (PC), etc. In the next section, we provide a theoretical proof that only four *f* –divergence based loss functions can be effectively utilized for the imputation of single-cell RNA sequencing data. The *f* -divergences are more general and computationally efficient, and can be used in different GAN architectures to offer a flexible and efficient way to measure the distance between probability distributions. Next, we will discuss our sc-*f* GAIN method, *f* -divergence based generative adversarial imputation network, for the imputation of missing values in the scRNA-seq data.

### sc-*f* GAIN: *f* -divergence based generative adversarial imputation network

The generative adversarial imputation nets (GAIN) method [34] was proposed to recover missing values implemented by a vanilla GAN, which uses binary cross-entropy (CE) loss to train models. Sometimes the cross-entropy divergence is not suitable for some complex distributions of scRNA-seq data. The sc-*f* GAIN algorithm is proposed to integrate *f* -divergence with GAIN model to build a more universal and efficient imputation architecture. Fig 1 illustrates the workflow of our sc-*f* GAIN method to impute missing values of scRNA-seq data, and the theoretical proof is provided in the next section.

**Fig 1.** An illustration of the scRNA-seq f-divergence based generative adversarial imputation network (sc-*f* GAIN) architecture. The generator takes as input the incomplete scRNA-seq data and a corresponding mask matrix, along with a random matrix to generate synthetic imputed data. The discriminator distinguishes between real and imputed values generated by the generator. Both generator and discriminator are trained using *f* -divergence loss functions.

### Generator and Discriminator

We adopt the notations used in Yoon *et al*.’s work [34] for clarity and coherence. The data vector **X** = (*X*_1_, …, *X*_*d*_) measures the expression levels of *d* genes and may contain both observed and missing values. To identify which components are missing or observed, a mask vector **M** = (*M*_1_, …, *M*_*d*_) is introduced, where *M*_*i*_ ∈ {0, 1} denotes whether the *i*-th component is observed (*M*_*i*_ = 1) or missing (*M*_*i*_ = 0).

One key component of the sc-*f* GAIN architecture is Generator *G*, which is implemented using convolutional neural networks. The Generator takes the sc-RNAseq dataset **X**, random noise vector **Z**, and binary mask **M** as inputs and generates imputed values for the missing observations. The input can be represented as the product of the element-wise multiplication between the mask matrix **M** and the raw data matrix **X**, and the element-wise multiplication between the complement of mask matrix, (**1** − **M**) and random noise matrix **Z**: **M** ⊙ **X** + (**1**− **M**) ⊙**Z**. The output of the Generator, denoted as *G*(**X, M, Z**), will be used to fill in the dropouts in the observed data **X** using the non-missing entries of **X** and the noise vector **Z**. The complete data, represented by 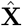, will be used as input to the discriminator. It comprises of the combination of imputed values *G*(**X, M, Z**) generated by *G* and the observed elements of **X**, which is expressed as

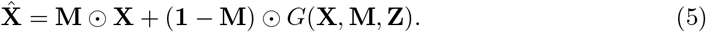

The second key component of the sc-*f* GAIN architecture is Discriminator *D*, which acts as an adversary or judge. Both the observed data and imputed values produced by the Generator network are input to the Discriminator network. The Discriminator’s role is to distinguish between real (observed) data and imputed data, thereby predict the binary mask vector **M**. Theorem 1-2 in the next section identified four *f* -divergence functionss (including CE, FKL, RKL, and JS), and proved that, given a fixed generator, there always exists one optimal discriminator; and given an optimal discriminator, we can always obtain optimal generators. In order to obtain a unique optimal solution for the Generator network *G*, Theorem 3 and Yoon’s work [34] have proved that the input to the Discriminator should not only consist of generated “imputed” values and observed data, but also some additional information, such as the hint matrix **H**, which should be designed to contain adequate information about the binary mask vector **M** in order to facilitate the training of the Discriminator *D*. Yoon *et al*’s work [34] introduced a binary random variable **B** = (*B*_1_,. .., *B*_*d*_) ∈ {0, 1}^*d*^, which indicates the location of missing values in the input data. To determine the value of *B*_*j*_ in the equation, a random value *k* is sampled uniformly from the set 1,. .., *d*, then, *B*_*j*_ = 1 if *j* ≠ *k*, and *B*_*j*_ = 0 otherwise. If **B** and **M** are independent, according to [34], hint matrix is expressed as:

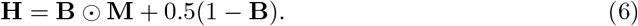

Theorem 4 proves that if the hint variable **H** is sampled using Eq. 6, the generator has the capability to replicate the desired distribution of the data, resulting in a unique distribution. The proof in Yoon *et al*’s work [34] assumes the binary cross-entropy adversarial loss. Our theoretical studies in the next section prove that this finding holds true if we use four different *f* -divergence functions for the adversarial loss.

### *f* -divergence based objective functions

The objective of sc-*f* GAIN is to impute missing values by training a generative model to learn to produce realistic data samples from some incomplete dataset. To achieve this, the objective functions, which serve as a measure of how well the generator and discriminator are performing their respective tasks, will be used to guide the optimization process towards better imputation results. Specifically, the generator network is trained to reduce the difference between the imputed data and real data, whereas the discriminator network is trained to differentiate between the imputed data and real data.

The input of discriminator includes the “complete” data 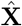 described by the Eq. 5, which combines both imputed values *G*(**X, M, Z**) generated by *G* and the observed elements of **X**, and hint matrix **H**. The output of the discriminator is 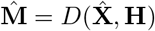, given the mask matrix **M**, the vanilla GAIN [34]‘s objective function for the discriminator *D* is described by a cross entropy function:

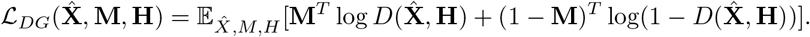

Now, we generalize the objective function of cross entropy to *f* -divergence, given the output of the network 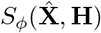 and output activation function *g*_*f*_, we can rewrite the Eq. 4 and get the *f* -divergence based objective function for the discriminator in the sc-*f* GAIN architecture:

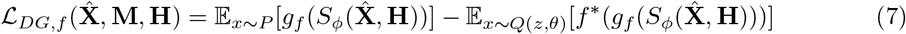

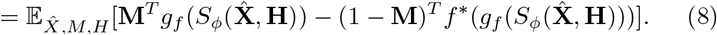

To train the discriminator *D*, we first fix the generator *G* and sample mini-batches of size *k*_*D*_ from the dataset, then, we optimize the discriminator by minimizing the loss function *ℒ*_*DG,f*_ over the mini-batches: 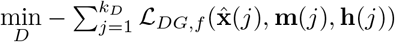. Table 1 summarizes five objective functions of discriminator based on different *f* –divergence functions and activation functions *g*_*f*_, where the sigmoid function 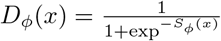 is applied on the output of the discriminator network *S*_*ϕ*_ (*x*). The abbreviations CE, FKL, RKL, JS, and PC will be used to represent the divergence of cross entropy, forward KL, reverse KL, Jensen-Shannon, and Pearson *χ*^2^ respectively.

**Table 1.**
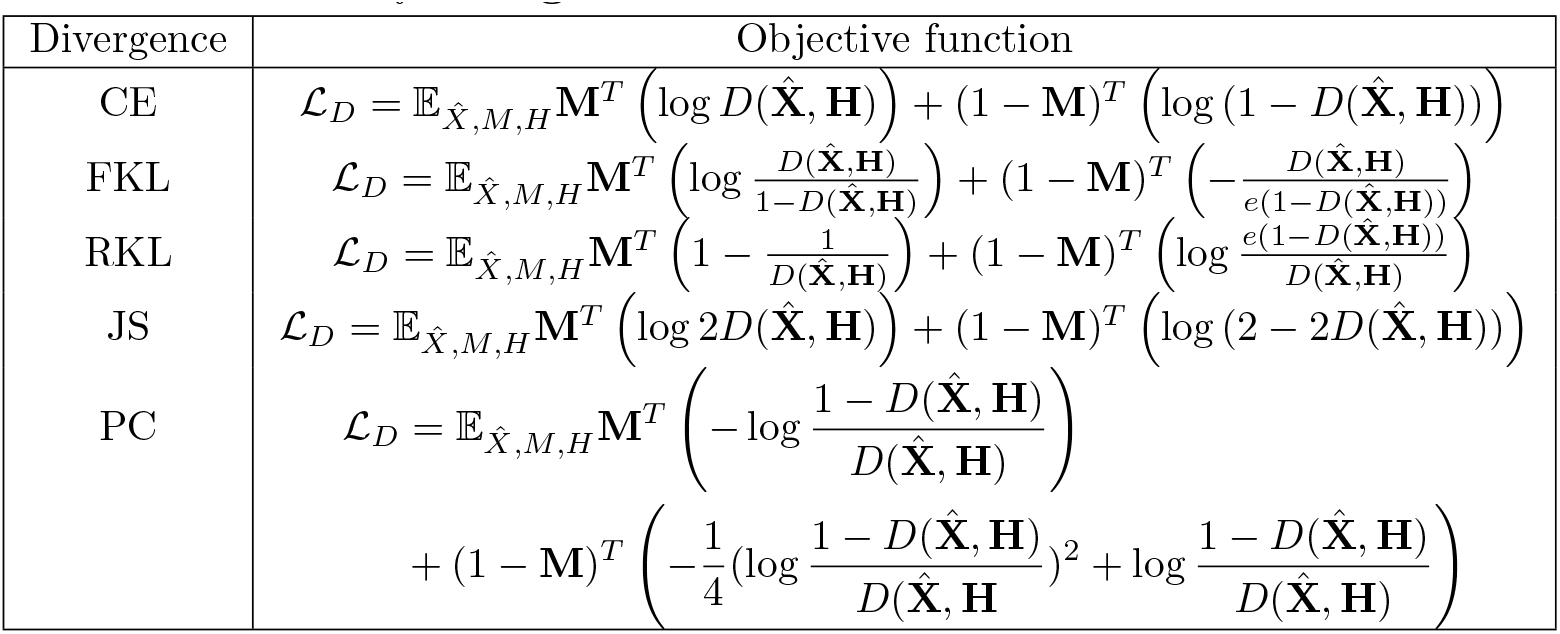
Objective functions of the Discriminator in the sc-*f* GAIN models based on different *f* -divergence function.

After the discriminator *D* is updated, the next step is to optimize the generator *G*. The generator’s optimization is dependent on two loss functions: reconstruction loss *ℒ*_*R*_ for the observed entries (*m* = 1) and adversarial loss ℒ_*GA,f*_ for the missing entries (*m* = 0). Similar to the original GAIN [34], we use the mean squared error (MSE) to calculate the sc-*f* GAIN’s reconstruction loss between the generated data and the original data: ℒ_*R*_(**X, X**^*′*^) = ‖ **X X′‖** ^2^, where, ‖· ‖ is the Euclidean norm, and **X**^*′*^ = *G*(**X, M, Z**) is imputed value matrix given the observed value matrix **X**. Minimizing the reconstruction loss ℒ_*R*_ ensures that the generated values for the observed entries (*m* = 1) are as close as possible to the original values. Minimizing the adversarial loss ℒ_*GA,f*_ will train the generator to generate “realistic” imputed values for the missing elements (*m* = 0) that are difficult for the discriminator to distinguish from real values. The *f* -divergence based adversarial loss for the generator ℒ_*GA,f*_ can be derived by setting **M** = 0 in the Eq. 8:

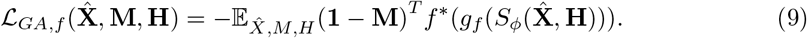

The overall objective function ℒ_*Gf*_ for the generator can be written as a weighted sum of these two loss functions, where the weight *λ* is a hyperparameter that need to be tuned to balance between the reconstruction accuracy and the data distribution fidelity for better imputation performance:

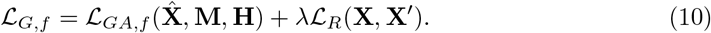

Algorithm 1 presents the pseudo-code for the sc-*f* GAIN method, which employs different *f* -divergences, instead of the cross entropy used in the original GAIN [34], to quantify the difference between two distributions. The sc-*f* GAIN imputation process involves training both generator and discriminator in order to minimize the *f* -divergence based loss functions defined by Eq. 8 and Eq. 10, respectively. The replacement of cross entropy in the loss function by the *f* -divergence is a non-trivial task since some divergence functions can not be used for the GAIN’s imputation procedure. In the next section, we will perform a theoretical analysis of our sc-*f* GAIN algorithm.

#### Algorithm 1 Pseudo-code of sc-*f* GAIN

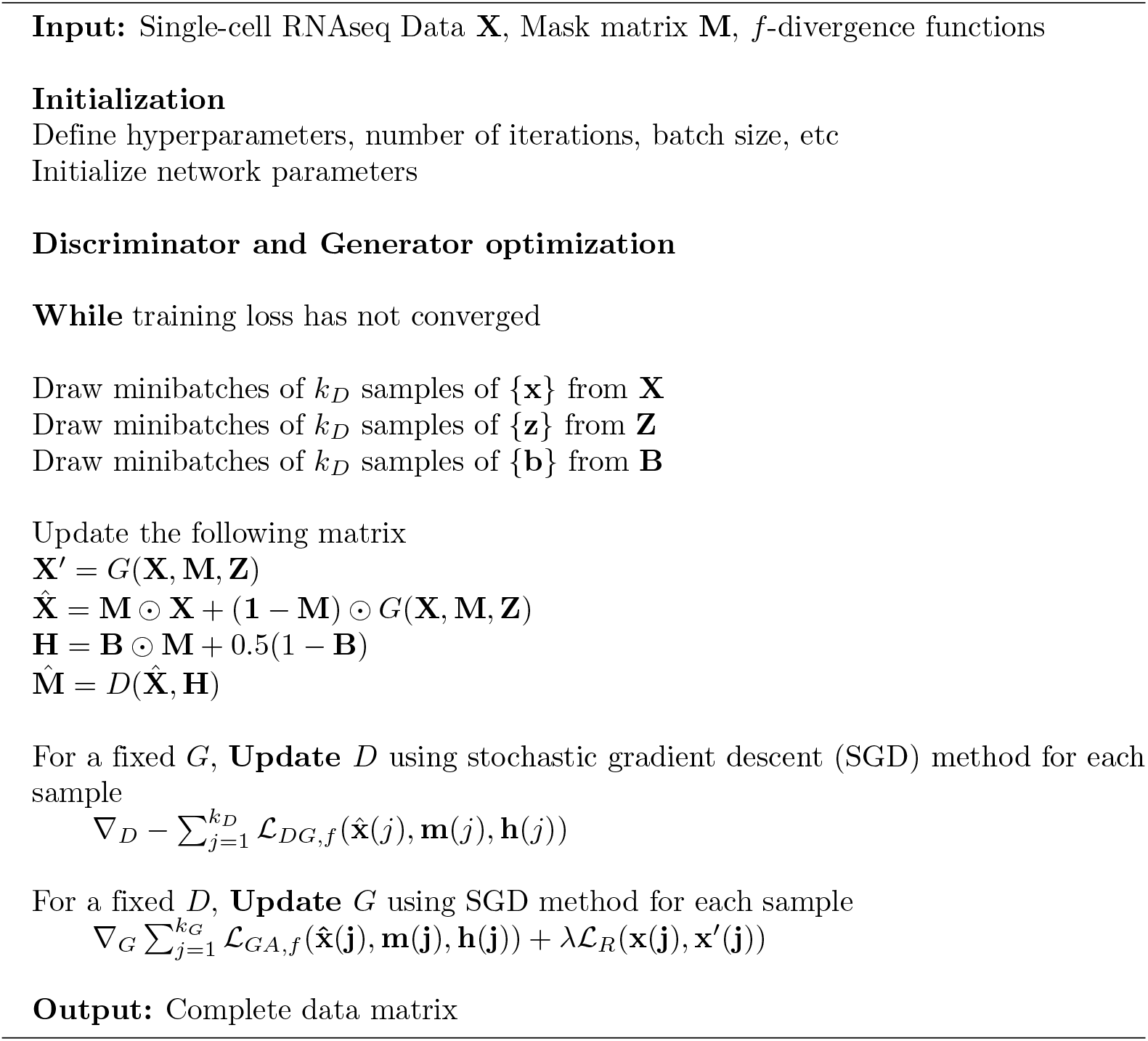

### Theoretical Analysis of sc-*f* GAIN Algorithm

In this section, we will identify specific *f* -divergence functions that can be used for the generative adversarial imputation network, and provide mathematical proof for the Algorithm 1. We adopt some notations and assumptions in Yoon *et al*’s work [34], and assume that **X** is independent of **M**, where *p*(**x, m, h**) denotes the joint distribution for the random variables 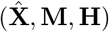, and 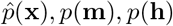 are corresponding marginal distributions.

#### Theorem 1.

*Let S*_*ϕ*_ (**x, h**) *be a function: χ* ⟶ *ℛ, where x*∈ χ, **h**∈ ℌ *(hint space), and p*(**x, h**) *>* 0, *D be a function: χ ⟶*[0, 1]^*d*^. *If the f-divergence based objective function is defined by the Eq. 8, then, given a fixed generator G, there always exists one optimal discriminator D*^*\**^(**x, h**) *if f* = *CE, FKL, RKL, JS, PC*.

*Proof*. The *f* -divergence based objective function Eq. 8 can be rewritten as

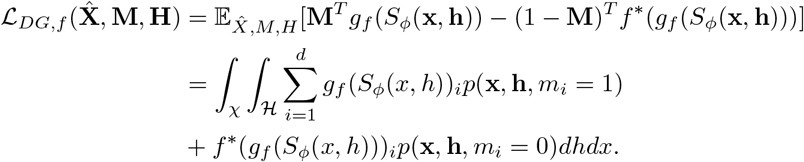

Given a fixed Generator *G*, the optimal Discriminator *D*^*\**^ is obtained by solving the equation 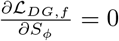, that is

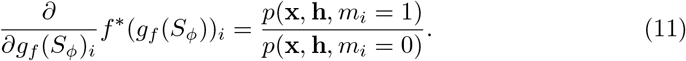

After inserting the *f* -divergence’s output activation functions and conjugate functions given in Table 2, and applying the sigmoid function 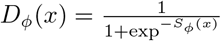 on the output of the discriminator network *S*_*ϕ*_ (*x*), we identified five *f* -divergences, including CE, FKL, RKL, JS, and PC, that always have an optimal discriminator *D*^*\**^ given a fixed *G*, for *i* ∈ {0, 1}^*d*^,

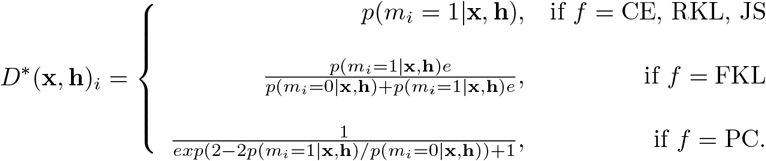

**Table 2.** *f* -divergence’s output activation function, conjugate function, and the optimal discriminator *D*^*\**^ for a given generator *G, p* = *p*(*x, h, m*_*i*_ = 1), and *q* = *p*(*x, h, m*_*i*_ = 0).

| $f$-Divergence | Output activation $g_f(s)$ | Conjugate $f^*(t)$ | Optimal $D^*$ |
| --- | --- | --- | --- |
| CE | $-\log(1 + \exp(-s))$ | $-\log(1 - \exp(t))$ | $\frac{p}{p+q}$ |
| FKL | $s$ | $\exp(t - 1)$ | $\frac{pe}{pe+q}$ |
| RKL | $-\exp(-s)$ | $-1 - \log(-t)$ | $\frac{p}{p+q}$ |
| JS | $\log(2) - \log(1 + \exp(-s))$ | $-\log(2 - \exp(t))$ | $\frac{p}{p+q}$ |
| PC | $s$ | $\frac{1}{4}t^2 + t$ | $\frac{\exp(2(p-q)/q)}{1+\exp(2(p-q)/q)}$ |

For a more detailed proof of Theorem 1, please refer to S1 Appendix.

If we substitute the optimal discriminator *D*^*\**^ derived in Theorem 1 into the objective function Eq. 8, we obtain the loss function of the generator *G* as follows:

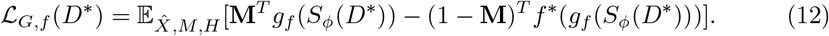

Then, by minimizing *ℒ*_*G,f*_ (*D*^*\**^), we derived the second theorem.

#### Theorem 2.

*The f-divergence based loss function ℒ*_*G,f*_ (*D*^*\**^) *has a global minimum if and only if the density p satisfies:*

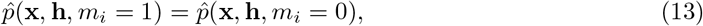

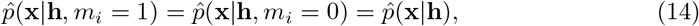

*for each i*∈ {1,. .., *d}, x* ∈ **X** *and ℌ such that p*(**h∣***m*_*i*_ = *t*) *>* 0. *And this theorem is true only if f* = *CE, FKL, RKL, JS*.

Yoon *et al*’s work [34] proved the validity of this theorem for the cross-entropy based loss function. We will prove that this theorem is also valid for the forward KL, reverse KL, and Jensen-Shannon divergence based loss functions described by Eq. 12, but it does not hold for the Pearson *∣*^2^ divergence.

*Proof*. We will present a concise proof of this theorem, focusing on the KL-divergence case, which is more intricate compared to the cross-entropy scenario. After substituting *D*^*\**^, using the Eq. 12 and objective function in the Table 1, the KL-divergence based loss function can be simplified as

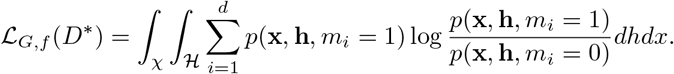

It follows that ℒ_*G,f*_ (*D*^*\**^) is minimized if and only if *p*(**x, h**, *m*_*i*_ = 1) = *p*(**x, h**, *m*_*i*_ = 0) for any *I* ∈ {1, …, *d*}.

The above loss function can also be rewritten as

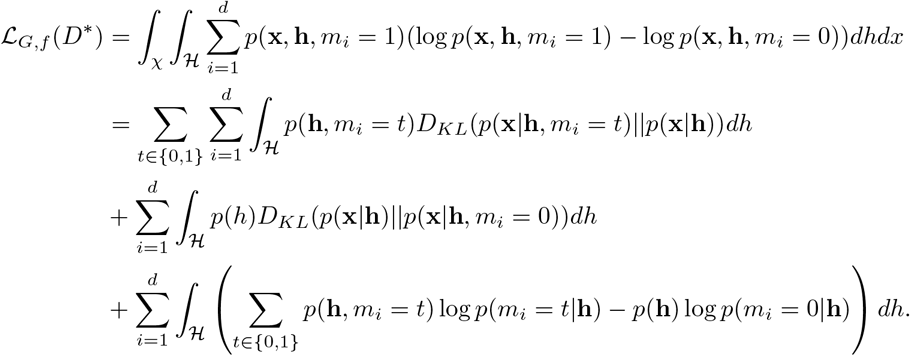

Since KL divergence *D*_*KL*_ is non-negative, so the loss function *ℒ*_*G,f*_ (*D*^*\**^) is minimized if and only if 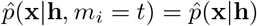 for any *i* ∈{1, …, *d*}. The detailed proof for different *f* -divergence cases are given in the S1 Appendix.

In comparison to [34], our work in Theorem 1-2 offers a more general proof based on the *f* -divergence functions, establishing that the optimal discriminator and generator can be attained using the sc-*f* GAIN algorithm when the loss function is formulated using four distinct *f* -divergence functions: cross-entropy, KL, reverse KL, and JS divergence. Theorem 2 demonstrates the independence of **x** from the mask variable **M** given the hint variable **H**. The amount of information contained in **H** directly influences the learning capability of the generator *G*. If **H** contains less informative hints or lacks important information, the learning ability of the generator may be compromised, which is discussed in the Theorem 3.

#### Theorem 3.

*In the sc-f GAIN algorithm, for f* = *CE, FKL, RKL, and JS, if the hint variable* **H** *is independent of mask variable* **M**, *then the density 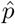 in the Theorem 2 is not unique*.

*Proof*. Theorem 2 has proved that, 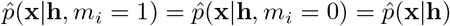 is valid for *f* = CE, FKL, RKL, and JS. If **H** is independent of **M**, and **H** is conditionally independent of **X** given **M**, it is easy to verify that 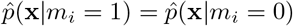, for all *I* ∈ {1, …, *d*}. Follow the same argumentation as [34] for the cross-entropy case, there are more parameters than the number of equations, so the density 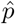 is not unique.

To get a unique density solution, a hinting mechanism is needed such that **H** reveals some information of **M** to the discriminator *D*, which means that they are not independent. In the last section, we adopt the method proposed in [34] to sample the hint variable using the Eq. 6, and assume **B** and **M** are independent. This hinting mechanism can ensure that the generator is capable of replicating the desired distribution of the data, that is the Theorem 4.

#### Theorem 4.

*If the hint variable* **H** *is sampled according to Eq. 6, then the density* 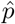 *in Theorem 2 is unique and satisfies* 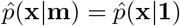*for any vector* **m**∈ 0, 1^*d*^ *and f* = *CE, FKL, RKL, JS, where* 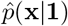 *is the density of* **X**. *That is, the distribution of imputed data is same as the distribution of original data*.

*Proof*. The proof is similar to the CE scenario [34]. Theorem 2 has shown that 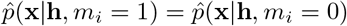 holds for the *f* -divergence of CE, FKL, RKL and JS. Because of Eq. 6, 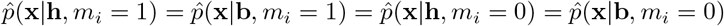 is valid. Since **B** and **M** are independent, it is easy to prove 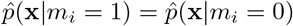. It means, for any two vectors **m**_**1**_, **m**_**2**_ ∈ *{*0, 1*}*^*d*^ that differ only on one component, we have 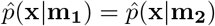.

This equation also holds true for any two vectors **m**_**1**_ and **m**_**2**_ in *{*0, 1*}*^*d*^, because we can always find a sequence of vectors between **m**_**1**_ and **m**_**2**_, such that all the adjacent vectors differ from each other in only one component. Consequently, the imputed data distribution 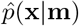 is the same for all possible vectors **m**∈ *{*0, 1} ^*d*^. This unique imputed data density, denoted by 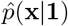, corresponds to the true data **X**’s density *p*(**x**), that is, 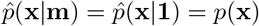. The proof is based on the Theorem 2, so it is true for *f* = CE, FKL, RKL, JS.

Theorem 1-4 theoretically confirm that the generative adversarial imputation network method remains valid if and only if the loss function is defined using four *f* -divergence, including CE, FKL, RKL, and JS divergence. The flexibility offered by the *f* -divergence formulation allows sc-*f* GAIN to accommodate various types of data and distributions, making it a more universal approach for imputing missing values.

## Results and Discussion

In this section, we will implement the sc-*f* GAIN algorithm to impute missing values in single-cell RNA sequencing data. The sc-*f* GAIN algorithm leverages *f* -divergence and generative adversarial networks to generate imputations that can capture the complex dependencies within the data without relying on any assumptions. To evaluate the effectiveness of the sc-*f* GAIN algorithm, we compared its performance with that of other state-of-the-art imputation methods including MAGIC [21], scImpute [14], and PBLR [23]. We designed several experiments to assess the performance of each method on the single-cell RNAseq data with varying missing rates, different metrics and setups. These experiments aim to determine the ability of each method to accurately impute missing values while preserving the underlying biological information.

### Data and Configuration

In our experiments, we evaluated the effectiveness of our sc-*f* GAIN algorithm and various imputation techniques using real single-cell RNAseq data from the UMI-based CellBench dataset GSE118767 [39]. This benchmark dataset, obtained through the 10x Chromium Genomics protocol [40], is widely used in current sc-RNAseq research and includes 3,918 cells comprising five human lung adenocarcinoma cell lines (H2228, H1975, A549, H838, and HCC827) equally mixed in the dataset. The dataset contains 10,164 genes, providing a comprehensive transcriptional profile of the mixed cell populations. In addition, we also utilized ten bulk RNA-seq samples from GSE86337 [41], which contains two replicates for each of the five cell lines. By using these two datasets, we were able to assess the performance of various imputation methods.

Before analyzing scRNA-seq gene expression data, data processing is performed, including data filtering and quality control as the first step. Due to the high dropout rate in single-cell RNAseq expression data, only genes that exhibit high differential expression in both raw single-cell RNA sequencing data GSE118767 and microarray bulk RNA sequencing data GSE86337 are retained. The data is then normalized by library size and subjected to log-transformation before being fitted into the model. The output is subsequently renormalized to ensure consistency.

Our simulation studies revealed that running the sc-*f* GAIN algorithm for 10,000 iterations provided sufficient training time for the generator and discriminator networks to converge to an optimal state. This convergence resulted in the generator producing more accurate imputations, while the discriminator effectively distinguished between real and generated data. Therefore, we consistently configured the model to run for 10,000 iterations in our experiments, varying the missing rate for the mask matrix. To ensure consistency with previous research, we adopted the same configuration as [34], while exploring different values for the hyperparameter of reconstruction loss.

Additionally, we set the hint rate, which is used to generate the hint matrix from the mask vector, to 0.9. Previous studies [34] have shown that this hint rate yields the best performance. The values of hyperparameters and sc-*f* GAIN code are released on the Github: https://github.com/TongSii/sc-fGAIN.

### Real Data Analysis

Next, we designed different experiments and used real scRNA-seq data with known ground truth values to evaluate the accuracy of sc-*f* GAIN’s imputation algorithm based on different *f* -divergence functions and compare with other imputation methods. In the first type of experiment, we intentionally introduced missing values into a portion of the data with varying rates of missingness to assess the effectiveness of sc-*f* GAIN in imputing missing values.

To validate the imputation accuracy of sc-*f* GAIN algorithm and other imputation methods, we computed the root-mean-square error (RMSE) between the true values and the imputed values in the scRNA-seq data, which has been used in many studies [34, 42]. The root mean squared error(RMSE) is defined as 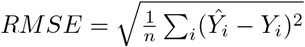, where *Y*_*i*_ and *Ŷ*_*i*_ represent true values and imputed values respectively.

#### sc-*f* GAIN can efficiently impute missing values

Most traditional imputation methods do not perform well when the cells are difficult to differentiate from one another. In order to achieve a high level of imputation accuracy for comparison purposes, we select a small subset of genes showing significant differential expression (DE) for the imputation process. Many methods [43, 44, 45], including p-value and fold-change method, have been used to identify differentially expressed genes. Similar to [43], we select the genes with p-values *<* 0.001 in both single-cell and bulk RNAseq data for the first round data analysis, and finally 269 highly differentially expressed genes were selected. Table 3 reports the performance metric RMSE for various imputation methods given different missing rates for the A549 cell line. The lower the RMSE score, the better the imputation performance of the method.

**Table 3.** Comparison of RMSE scores for the imputation of 269 highly differentially expressed genes of the A549 cell line.

| Method \ Missing rate | 0.1 | 0.2 | 0.3 | 0.4 | 0.5 |
| --- | --- | --- | --- | --- | --- |
| **MAGIC** | 0.14219 | 0.16442 | 0.18861 | 0.21870 | 0.24963 |
| **scImpute** | 0.22818 | 0.23586 | 0.25137 | 0.26979 | 0.28257 |
| **PBLR** | 0.32017 | 0.32251 | 0.32208 | 0.32192 | 0.32085 |
| **sc-*f*GAIN(CE)** | 0.17808 | 0.18376 | 0.17894 | 0.17297 | 0.18989 |
| **sc-*f*GAIN(FKL)** | 0.20592 | 0.21683 | 0.20993 | 0.21219 | 0.20616 |
| **sc-*f*GAIN(RKL)** | 0.21701 | 0.21042 | 0.21238 | 0.20262 | 0.20425 |
| **sc-*f*GAIN(JS)** | 0.17589 | 0.18052 | 0.17499 | 0.17978 | 0.18176 |

Table 3’s results show that, the sc-*f* GAIN method outperformed most traditional imputation methods, with the *CE* and *JS* based sc-*f* GAIN method achieving the smallest RMSE score. Our sc-*f* GAIN method demonstrates robustness and low sensitivity to the missing rates, and it outperforms all the traditional methods when the missing rate is very high. MAGIC only performed well when the missing rate was relatively low, but its performance deteriorated with higher dropout rates, moreover, PBLR and scImpute showed poor performance across all missing rates, and as the missing rate increases, all traditional methods’ imputation accuracy declines.

Furthermore, the results presented in Table 3 demonstrate that the performance of our sc-*f* GAIN algorithm is influenced by the selection of *f* -divergence functions. One major reason for the poor performance of traditional methods is the assumption that the missing values follow some specific distributions, which could not accurately capture the intricacy of the high-dimensional scRNA-seq data, and they rely on additional information about adjacent genes for constructing k-nearest neighbor graphs and clustering before imputation. Given that single-cell RNA sequencing data often exhibits a high dropout rate, our sc-*f* GAIN method outperforms many traditional methods and proves to be superior in handling scRNA-seq data. Table 4 presents the RMSE results for all cell lines. These results are consistent with the results reported for the single cell line in Table 3, that is, our sc-*f* GAIN method outperforms all the traditional methods when the missing rate is very high.

**Table 4.**
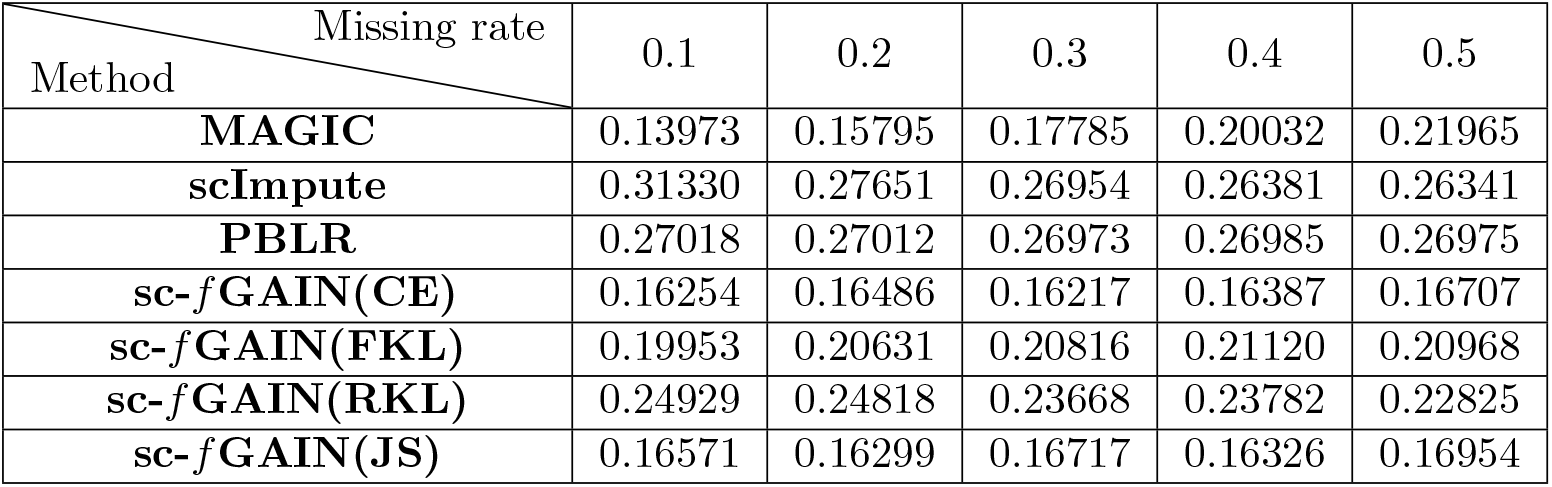
Comparison of RMSE scores for the imputation of 269 highly differentially expressed genes of all cell lines.

| Method \ Missing rate | 0.1 | 0.2 | 0.3 | 0.4 | 0.5 |
| --- | --- | --- | --- | --- | --- |
| **MAGIC** | 0.13973 | 0.15795 | 0.17785 | 0.20032 | 0.21965 |
| **scImpute** | 0.31330 | 0.27651 | 0.26954 | 0.26381 | 0.26341 |
| **PBLR** | 0.27018 | 0.27012 | 0.26973 | 0.26985 | 0.26975 |
| **sc-*f*GAIN(CE)** | 0.16254 | 0.16486 | 0.16217 | 0.16387 | 0.16707 |
| **sc-*f*GAIN(FKL)** | 0.19953 | 0.20631 | 0.20816 | 0.21120 | 0.20968 |
| **sc-*f*GAIN(RKL)** | 0.24929 | 0.24818 | 0.23668 | 0.23782 | 0.22825 |
| **sc-*f*GAIN(JS)** | 0.16571 | 0.16299 | 0.16717 | 0.16326 | 0.16954 |

To investigate the impact of data size on the performance of the sc-*f* GAIN algorithm, we divided the entire dataset across genes into three equally sized parts and conducted imputation experiments on each split dataset. We also calculated the RMSE metric between the observed dataset and the combined imputed split data, which allows us to compare the results with those presented in Table 3-4. The results in Table 5 demonstrate that the performance of the sc-*f* GAIN method is not very sensitive to the size of the data when the missing rate is low, but its performance will improve when the rate is high. Moreover, we observed that the sc-*f* GAIN method takes shorter running times to impute smaller split datasets. This observation presents an opportunity to leverage parallel computing and reduce imputation time by partitioning large datasets into smaller subsets across genes.

**Table 5.** RMSE score for the imputation of 269 DE genes on the split data.

| Method \ Missing rate | 0.1 | 0.2 | 0.3 | 0.4 | 0.5 |
| --- | --- | --- | --- | --- | --- |
| **Results for all five cell lines** |  |  |  |  |  |
| **sc-*f*GAIN(CE)** | 0.13838 | 0.14969 | 0.15151 | 0.15596 | 0.16318 |
| **sc-*f*GAIN(FKL)** | 0.20000 | 0.19921 | 0.20915 | 0.20619 | 0.20478 |
| **sc-*f*GAIN(RKL)** | 0.22684 | 0.25093 | 0.23430 | 0.23482 | 0.22825 |
| **sc-*f*GAIN(JS)** | 0.13346 | 0.14648 | 0.14698 | 0.15795 | 0.15761 |
| **Results for the A549 cell line** |  |  |  |  |  |
| **sc-*f*GAIN(CE)** | 0.11531 | 0.11917 | 0.12046 | 0.12025 | 0.12100 |
| **sc-*f*GAIN(FKL)** | 0.20050 | 0.21531 | 0.20917 | 0.20916 | 0.20836 |
| **sc-*f*GAIN(RKL)** | 0.21157 | 0.22077 | 0.21049 | 0.20686 | 0.20870 |
| **sc-*f*GAIN(JS)** | 0.11434 | 0.11850 | 0.11687 | 0.11714 | 0.12008 |

In another experiment, we included a larger dataset consisting of 2000 highly differentially expressed genes as input. These genes were selected using a similar approach as the work [25, 42]. The results in Table 6 align with those obtained from the small dataset presented in Table 3-4. Both sc-*f* GAIN (FKL) and MAGIC methods achieves the lowest RMSE scores across all missing rates, and the other two sc-*f* GAIN methods based on CE and JS divergences remain competitive compared with the PBLR and scImpute, particularly when dealing with the data with high dropout rates. Our sc-*f* GAIN method’s performance is influenced by the choice of loss function which is described by the *f* -divergence, as evident from the results in Tables 3-6. This indicates that sc-*f* GAIN offers a more comprehensive and versatile approach compared to the original GAIN model. To further explore the capabilities of sc-*f* GAIN, we have developed a convenient Huggingface tool. This tool, accessible at [46], provides users with the ability to identify the optimal *f* -divergence function for imputing a given dataset, leading to the lowest RMSE score. The corresponding code is hosted on GitHub at [47].

**Table 6.** Comparison of RMSE score (2000 genes) in the A549 cell line.

| Method \ Missing rate | 0.1 | 0.2 | 0.3 | 0.4 | 0.5 |
| --- | --- | --- | --- | --- | --- |
| **MAGIC** | 0.10309 | 0.09844 | 0.10168 | 0.11060 | 0.12283 |
| **scImpute** | 0.15627 | 0.15312 | 0.15401 | 0.15424 | 0.15479 |
| **PBLR** | 0.10545 | 0.13518 | 0.15233 | 0.16398 | 0.17242 |
| **sc-*f*GAIN(CE)** | 0.15206 | 0.15113 | 0.14939 | 0.15550 | 0.14953 |
| **sc-*f*GAIN(FKL)** | 0.12846 | 0.11287 | 0.11960 | 0.13058 | 0.12606 |
| **sc-*f*GAIN(RKL)** | 0.18133 | 0.17231 | 0.14557 | 0.14608 | 0.17078 |
| **sc-*f*GAIN(JS)** | 0.14958 | 0.15509 | 0.15329 | 0.15407 | 0.14922 |

Single-cell RNA sequencing data frequently encounters high dropout events, some traditional imputation methods make assumptions about the data distribution when filling in missing values. However, these assumptions can introduce bias and potentially lead to incorrect estimates of gene expression levels, thereby affecting downstream analysis. In contrast, the proposed sc-*f* GAIN method is a generative model that does not rely on any predefined data distribution assumptions. As a result, it is theoretically expected to mitigate the imputation bias compared to some traditional methods.

#### sc-*f* GAIN can reduce imputation bias

To assess the effectiveness of the imputation methods in reducing prediction bias, the gene-specific standard deviation across cells can be compared before and after imputation. Typically, successful imputation methods should result in a reduction of bias, indicated by a decrease in the standard deviation of gene expression levels after imputation compared to that without imputation.

Fig 2 demonstrates the changes of the standard deviation (SD) across different imputation methods in each cell line. Prior to imputation (red box), the SD values range from 0.1 to 1.4, with a median value exceeding 0.5. However, when applying the traditional imputation methods such as scImpute (brown box) and PBLR (green box), there is only a slight reduction in the SD values, which range from 0.1 to 1.0 with a median value around 0.3-0.5. It is important to note that a higher standard deviation indicates higher uncertainty or variability in the imputed values, reflecting the limitations of these two imputation methods. The relatively small reduction in SD values suggests that scImpute and PBLR are not able to effectively address the uncertainty and variability associated with missing values in the dataset. The poor performance of PBLR can be attributed to the assumption made regarding the prior distribution of the missing values, which may not accurately represent the true underlying distribution of the missing values. On the other hand, scImpute’s high prediction bias can be attributed to its assumption that the underlying gene expression data matrix has a low-rank structure, that is, the majority of the genetic information can be captured by a smaller number of latent factors. However, the scRNA-seq data exhibit more intricate patterns and dependencies that might not be adequately represented by a low-rank structure. These assumptions limit the ability of PBLR and scImpute to capture the complexity of the data, leading to a biased imputation.

**Fig 2.** Comparison of sc-*f* GAIN imputation methods and traditional methods in terms of gene-specific standard deviation. The x-axis represents different imputation methods, and y-axis represents the gene-specific standard deviation.

In contrast, our sc-*f* GAIN methods, by leveraging generative adversarial networks and utilizing different *f* -divergences (CE, FKL, RKL, JS), overcome the limitations imposed by the assumptions made in traditional imputation methods. When using the sc-*f* GAIN methods, the resulting standard deviation values are consistently lower compared to traditional imputation methods. All the median values are lower than 0.1 with small variations, which indicates a significant decrease in the variability of the imputed values compared to the initial missing values. Among the traditional methods, only MAGIC demonstrates comparable performance to our sc-*f* GAIN method. However, it is important to note that our sc-*f* GAIN method offers flexibility in choosing different *f*-divergences, allowing for better adaptation to diverse data patterns. This finding suggests that imputation using sc-*f* GAIN introduces less bias into the data compared to traditional methods. Consequently, sc-*f* GAIN enables accurate imputation of missing expression values without significantly altering the overall variability of the data, thereby facilitating improved downstream analyses of single-cell RNA sequencing data.

## Conclusion

In this work, we have developed a novel *f* -divergence based generative adversarial imputation network method, called sc-*f* GAIN, to impute missing values in single-cell RNA sequencing data. Unlike some traditional imputation methods, sc-*f* GAIN employs a generative model to generate synthetic values without making any distribution assumptions, the missing values are imputed by training two neural networks whose objective functions are described by *f* -divergence. Our theoretical studies, for the first time, identified only four *f* -divergences, namely cross-entropy, forward Kullback-Leibler (KL), reverse KL, and Jensen-Shannon, that can be effectively used as loss functions to train the sc-*f* GAIN algorithm to generate imputed values without any assumptions, and mathematically proved that the distribution of imputed data using sc-*f* GAIN algorithm is same as the distribution of original data.

Our experiments with public scRNA-seq data have shown that the proposed sc-*f* GAIN method has many advantages compared with some traditional imputation methods and vanilla GAIN. Specifically, the imputed values generated by sc-*f* GAIN have a smaller root-mean-square error compared to many traditional methods. Importantly, the performance of sc-*f* GAIN is robust to varying missing rates of scRNA-seq data, whereas most traditional methods’ performance will deteriorate with higher missing rates. The studies also indicate that the performance of sc-*f* GAIN is sensitive to the choice of *f* -divergence used in the loss function. Our tool can also help to identify a suitable *f* -divergence function to optimize the performance of the sc-*f* GAIN algorithm. Moreover, our investigation also revealed that the sc-*f* GAIN method can reduce imputation bias by producing smaller standard deviations of gene expression values compared to most traditional methods. This indicates that sc-*f* GAIN is capable of accurately imputing missing values without altering the overall variability of the data. Such an ability can help improve downstream analyses that rely on accurate estimation of gene expression levels.

Despite its promising potential, sc-*f* GAIN has some limitations that should be acknowledged. One such limitation is its training speed, particularly when working with large datasets. For instance, analyzing the scRNAseq dataset GSE118767 using sc-*f* GAIN can take approximately 3 hours on a Tesla V100-SXM2 16GB GPU. However, there are several approaches that can be implemented to mitigate this issue, including reducing the batch size, minimizing the number of iterations, and partitioning the dataset into smaller subsets for separate imputations. Our investigations have revealed that employing these strategies does not significantly compromise the accuracy of the results, making them feasible solutions for addressing the training speed limitation. In conclusion, our findings highlight the promising potential of sc-*f* GAIN in enhancing the quality of single-cell RNA sequencing data and facilitating more precise and reliable downstream analyses.

## Supporting information

Proof for the Theorem 1-2

## Supporting Information

**S1 Appendix. Proof of Theorem 1-2**. Detailed proof of the Theorem 1-2 in the Section: Theoretical analysis of sc-*f* GAIN Algorithm.

## Data Availability

The authors confirm that all data underlying the findings are fully available without restriction. The data that we analyzed can be obtained from the National Center for Biotechnology Information Gene Expression Omnibus (GEO) repository under the accession number GSE118767 and GSE86337. The sc-*f* GAIN code is released on the Github: https://github.com/TongSii/sc-fGAIN.

## Acknowledgements

The computations were performed using the HPC facility offered by ITS, the Saint Louis University.

## Funding

This work was partially supported by research funds from the National Institutes of Health (NIH) grant number R15GM148915 to HG, and intramural President’s Research Funds to HG. The content is solely the responsibility of the authors and does not necessarily represent the official views of the National Institutes of Health. The funders had no role in study design, data collection and analysis, decision to publish, or preparation of the manuscript.

## Author Contributions

TS designed the project, implemented the algorithm, performed analysis and interpretation of the data, wrote the manuscript. ZH implemented the algorithm and data analysis. JY implemented the algorithm. JH participated in the study design, supervision of the work and writing the manuscript. HG conceived the idea and initiated the study, and wrote the manuscript and supervised the overall work.

## Competing Interests

The authors have declared that no competing interests exist.

## Notes

### Competing Interest Statement

The authors have declared no competing interest.

