## Supplementary material for "A novel *f*-divergence based generative adversarial imputation method for scRNA-seq data analysis": Proof for the Theorem 1-2

### S1 Appendix: Proof of Theorem 1-2

#### Proof of Theorem 1

We will use the following equation,

$$\frac{\partial}{\partial g_f(S_\phi)_i} f^*(g_f(S_\phi))_i = \frac{p(\mathbf{x}, \mathbf{h}, m_i = 1)}{p(\mathbf{x}, \mathbf{h}, m_i = 0)}. \tag{1}$$

and Table 2 to prove the Theorem 1.
Set $p_i = p(\mathbf{x}, \mathbf{h}, m_i = 1)$, $q_i = p(\mathbf{x}, \mathbf{h}, m_i = 0)$, apply the Sigmoid function on the output of the discriminator network $S_\phi(x, h)$, that is $D_i = \frac{1}{1+\exp(-S_\phi(x,h)_i)}$.

Below, we will derive the optimal discriminator $D_i^*$ given different $f$-divergence functions:

- $f$ = **Cross-entropy (CE):**

  $$\frac{\partial}{\partial g_f(S_\phi)_i} f^*(g_f(S_\phi))_i = \frac{p_i}{q_i} = \frac{\exp(-\log(1 + \exp(-S_\phi(x,h)_i)}{1 - \exp(-\log(1 + \exp(-S_\phi(x,h)_i)} = \frac{\frac{1}{1+\exp(-S_\phi(x,h)_i)}}{1 - \frac{1}{1+\exp(-S_\phi(x,h)_i)}}$$

  The above equation can be simplified as $\frac{p_i}{q_i} = \frac{D_i}{1-D_i}$.
  Finally, we get the optimal discriminator $D_i^* = \frac{p_i}{p_i+q_i}$.

- $f$ = **Kullback-Leibler(FKL)**
  $\frac{\partial}{\partial g_f(S_\phi)_i} f^*(g_f(S_\phi))_i = \frac{p_i}{q_i} = \exp(-\log(\frac{1}{S_\phi(x,h)_i} - 1) - 1)$
  The above equation can be simplified as $\frac{p_i}{q_i} = \frac{D_i}{e(1-D_i)}$.
  Finally, we get the optimal discriminator $D_i^* = \frac{p_i e}{p_i e + q_i}$.

- $f$ = **Reverse Kullback-Leibler (RKL)**
  $\frac{\partial}{\partial g_f(S_\phi)_i} f^*(g_f(S_\phi))_i = \frac{p_i}{q_i} = \frac{1}{-\exp(-S_\phi(x,h)_i)}$
  The above equation can be simplified as $\frac{p_i}{q_i} = \frac{D_i}{1-D_i}$.
  Finally, we get the optimal discriminator $D_i^* = \frac{p_i}{p_i+q_i}$

- $f$ = **Jensen-Shannon (JS)**
  $\frac{\partial}{\partial g_f(S_\phi)_i} f^*(g_f(S_\phi))_i = \frac{p_i}{q_i} = \frac{\frac{2}{1+\exp(-S_\phi(x,h)_i)}}{2 - \frac{2}{1+\exp(-S_\phi(x,h)_i)}}$
  The above equation can be simplified as $\frac{p_i}{q_i} = \frac{D_i}{1-D_i}$.
  Finally, we get the optimal discriminator $D_i^* = \frac{p_i}{p_i+q_i}$.

- $f$ = **Pearson $\chi^2$ (PC)**
  $\frac{\partial}{\partial g_f(S_\phi)_i} f^*(g_f(S_\phi))_i = \frac{p}{q} = \frac{1}{2} S_\phi(x,h)_i + 1$
  Finally, we get the optimal discriminator $D_i^* = \frac{exp(2(p_i - q_i)/q_i)}{1+exp(2(p_i-q_i)/q_i)}$.

#### Proof of Theorem 2

Given the optimal discriminator $D_i^*$, for each $f$-divergence function, the objective function $\mathcal{L}_{G,f}(D^*)$ can be expressed as:

- $f$ **= CE**

$$\mathcal{L}_{G,f}(D^*) = \sum_{i}^{d} \int_{\chi} \int_{\mathcal{H}} \left(p_i \log(D_i^*) + q_i \log(1 - D_i^*)\right) dxdh$$
$$= \sum_{i}^{d} \int_{\chi} \int_{\mathcal{H}} \left(p_i \log(\frac{p_i}{p_i + q_i}) + q_i \log(\frac{q_i}{p_i + q_i})\right) dxdh$$

It can also be written as:

$$\mathcal{L}_{G,f}(D^*) = \sum_{t\in\{0,1\}} \sum_{i=1}^{d} \int_{\chi} \int_{\mathcal{H}} p(\mathbf{x}, \mathbf{h}, m_i = t) \log(\frac{p(\mathbf{x}, \mathbf{h}, m_i = t)}{p(\mathbf{x}, \mathbf{h}, m_i = 0) + p(\mathbf{x}, \mathbf{h}, m_i = 1)}) dhdx$$
$$= \sum_{t\in\{0,1\}} \sum_{i=1}^{d} \int_{\chi} \int_{\mathcal{H}} p(\mathbf{x}, \mathbf{h}, m_i = t) \log(\frac{p(\mathbf{x}, m_i = t|\mathbf{h})p(m_i = t|h)}{p(\mathbf{x}|\mathbf{h})p(m_i = t|h)}) dhdx$$
$$= \sum_{t\in\{0,1\}} \sum_{i=1}^{d} \int_{\chi} \int_{\mathcal{H}} p(\mathbf{h}, m_i = t)p(\mathbf{x}|\mathbf{h}, m_i = t) \log(\frac{p(\mathbf{x}|\mathbf{h}, m_i = t)}{p(\mathbf{x}|\mathbf{h}))} dhdx$$
$$+ \sum_{t\in\{0,1\}} \sum_{i=1}^{d} \int_{\mathcal{H}} p(\mathbf{h}, m_i = t) \log(p(m_i = t|\mathbf{h}))dh$$
$$= \sum_{t\in\{0,1\}} \sum_{i=1}^{d} \int_{\mathcal{H}} p(\mathbf{h}, m_i = t) D_{KL}(p(x|\mathbf{h}, m_i = t)||p(x|\mathbf{h}))dh$$
$$+ \sum_{t\in\{0,1\}} \sum_{i=1}^{d} \int_{\mathcal{H}} p(\mathbf{h}, m_i = t) \log(p(m_i = t|\mathbf{h}))dh$$

Since KL divergence is non-negative, so the loss function $\mathcal{L}_{G,f}(D^*)$ is minimized if and only if $\hat{p}(\mathbf{x}|\mathbf{h}, m_i = t) = \hat{p}(\mathbf{x}|\mathbf{h})$ for any $i \in \{1, ..., d\}$.

- $f$ = **FKL**

$$
\begin{aligned}
\mathcal{L}_{G,f}(D^*) &= \sum_{i=1}^{d}\int_{\chi}\int_{\mathcal{H}}\left(-p(\frac{1}{D^*}-1)+q(1+\log(\frac{1}{D^*}-1))\right)dxdh \\
&= \sum_{i=1}^{d}\int_{\chi}\int_{\mathcal{H}}\left(p\log\frac{p}{q}\right)dxdh \\
&= \sum_{i=1}^{d}\int_{\chi}\int_{\mathcal{H}}\left(p\log p+q\log q-(p+q)\log q\right)dxdh \\
&= \sum_{i=1}^{d}\int_{\chi}\int_{\mathcal{H}}\sum_{t\in\{0,1\}}p(\mathbf{x},\mathbf{h},m_i=t)\log\frac{P(x|h,m_i=t)}{P(x|h)}dxdh \\
&+ \sum_{t\in\{0,1\}}\sum_{i=1}^{d}\int_{\mathcal{H}}p(\mathbf{h},m_i=t)\log(p(m_i=t|\mathbf{h}))dh \\
&- \sum_{i=1}^{d}\int_{\chi}\int_{\mathcal{H}}p(x,h)\log p(m_i=0|x,h)dxdh \\
&= \sum_{i=1}^{d}\int_{\chi}\int_{\mathcal{H}}\sum_{t\in\{0,1\}}p(\mathbf{x}|\mathbf{h},m_i=t)p(\mathbf{h},m_i=t)\log\frac{P(x|h,m_i=t)}{P(x|h)}dxdh \\
&+ \sum_{t\in\{0,1\}}\sum_{i=1}^{d}\int_{\mathcal{H}}p(\mathbf{h},m_i=t)\log(p(m_i=t|\mathbf{h}))dh \\
&- \sum_{i=1}^{d}\int_{\chi}\int_{\mathcal{H}}p(h)p(x|h)\log\frac{p(x,m_i=0|h)}{p(x|h)}dxdh \\
&= \sum_{t\in\{0,1\}}\sum_{i=1}^{d}\int_{\mathcal{H}}p(\mathbf{h},m_i=t)D_{KL}(p(x|\mathbf{h},m_i=t)||p(x|\mathbf{h})) \\
&+ p(\mathbf{h},m_i=t)\log(p_m(m_i=t|\mathbf{h}))dh \\
&+ \sum_{i=1}^{d}\int_{\mathcal{H}}p(h)D_{KL}(p(x|h)||p(x|m_i=0,h))-p(h)\log p(m_i=0|h)dh
\end{aligned}
$$

Similar to CE's proof, since KL divergence is non-negative, so $\mathcal{L}_{G,f}(D^*)$ is minimized if and only if $p(x|\mathbf{h},m_i=t)=p(x|\mathbf{h})$ for the 1st and 2nd term.

- $f$ **= RKL**

$$
\begin{aligned}
\mathcal{L}_{G,f}(D^*) &= \sum_{i=1}^{d}\int_{\chi}\int_{\mathcal{H}}\left(p\log(\frac{D^*}{1-D^*}) - \frac{q}{e}\frac{D^*}{1-D^*}\right)dxdh \\
&= \sum_{i=1}^{d}\int_{\chi}\int_{\mathcal{H}}\left(p\log(\frac{pe}{q}) - \frac{q}{e}\log(\frac{pe}{q})\right)dxdh = \sum_{i=1}^{d}\int_{\chi}\int_{\mathcal{H}}\left(q\log\frac{q}{p}\right)dxdh \\
&= \sum_{i=1}^{d}\int_{\chi}\int_{\mathcal{H}}\left(q\log q + p\log p - (p+q)\log(p)\right)dxdh \\
&= \sum_{i=1}^{d}\int_{\chi}\int_{\mathcal{H}}\sum_{t\in\{0,1\}} p(\mathbf{x},\mathbf{h},m_i=t)\log\frac{P(x|h,m_i=t)}{P(x|h)}dxdh \\
&+ \sum_{t\in\{0,1\}}\sum_{i=1}^{d}\int_{\mathcal{H}} p(\mathbf{h},m_i=t)\log(p(m_i=t|\mathbf{h}))dh \\
&- \sum_{i=1}^{d}\int_{\chi}\int_{\mathcal{H}} p(x,h)\log p(m_i=1|x,h)dxdh \\
&= \sum_{i=1}^{d}\int_{\chi}\int_{\mathcal{H}}\sum_{t\in\{0,1\}} p(\mathbf{x}|\mathbf{h},m_i=t)p(\mathbf{h},m_i=t)\log\frac{P(x|h,m_i=t)}{P(x|h)}dxdh \\
&+ \sum_{t\in\{0,1\}}\sum_{i=1}^{d}\int_{\mathcal{H}} p(\mathbf{h},m_i=t)\log(p(m_i=t|\mathbf{h}))dh \\
&- \sum_{i=1}^{d}\int_{\chi}\int_{\mathcal{H}} p(h)p(x|h)\log\frac{p(x,m_i=1|h)}{p(x|h)}dxdh \\
&= \sum_{t\in\{0,1\}}\sum_{i=1}^{d}\int_{\mathcal{H}} p(\mathbf{h},m_i=t)D_{KL}(p(x|\mathbf{h},m_i=t)||p(x|\mathbf{h})) \\
&+ p(\mathbf{h},m_i=t)\log(p_m(m_i=t|\mathbf{h}))dh \\
&+ \sum_{i=1}^{d}\int_{\mathcal{H}} p(h)D_{KL}(p(x|h)||p(x|m_i=1,h)) - p(h)\log p(m_i=1|h)dh
\end{aligned}
$$

Similarly to FKL, since KL divergence is non-negative, so, to get $\mathcal{L}_{G,f}(D^*)$ minimized if and only if $p(x|\mathbf{h},m_i=t)=p(x|\mathbf{h})$.

- $f$ = **JS**

$$
\begin{aligned}
\mathcal{L}_{G,f}(D^*) &= \sum_{t\in\{0,1\}}\sum_{i=1}^{d}\int_{\mathcal{X}}\int_{\mathcal{H}} \log(2D^*) + q\log(2-2D^*) \\
&= \sum_{t\in\{0,1\}}\sum_{i=1}^{d}\int_{\mathcal{X}}\int_{\mathcal{H}} p\log\,(\frac{2p}{p+q}) + q\log\,(\frac{2q}{p+q}) \\
&= \sum_{t\in\{0,1\}}\sum_{i=1}^{d}\int_{\mathcal{X}}\int_{\mathcal{H}} p(\mathbf{x},\mathbf{h},m_i=t)\log(\frac{p(\mathbf{x},\mathbf{h},m_i=t)}{p(\mathbf{x},\mathbf{h},m_i=0)+p(\mathbf{x},\mathbf{h},m_i=1)})dhdx \\
&+ \sum_{t\in\{0,1\}}\sum_{i=1}^{d}\int_{\mathcal{X}}\int_{\mathcal{H}} p(\mathbf{x},\mathbf{h},m_i=t)\log\,(2)dxdh \\
&= \sum_{t\in\{0,1\}}\sum_{i=1}^{d}\int_{\mathcal{X}}\int_{\mathcal{H}} p(\mathbf{x},\mathbf{h},m_i=t)\log(\frac{p(\mathbf{x},m_i=t|\mathbf{h})p_m(m_i=t|h)}{p(\mathbf{x}|\mathbf{h})p_m(m_i=t|h)})dhdx + C \\
&= \sum_{t\in\{0,1\}}\sum_{i=1}^{d}\int_{\mathcal{X}}\int_{\mathcal{H}} p(\mathbf{h},m_i=t)p(\mathbf{x}|\mathbf{h},m_i=t)\log(\frac{p(\mathbf{x}|\mathbf{h},m_i=t)}{p(\mathbf{x}|\mathbf{h}))}dhdx \\
&+ \sum_{t\in\{0,1\}}\sum_{i=1}^{d}\int_{\mathcal{H}} p(\mathbf{h},m_i=t)p(\mathbf{x}|\mathbf{h},m_i=t)\log(p_m(m_i=t|\mathbf{h}))dh + C \\
&= \sum_{t\in\{0,1\}}\sum_{i=1}^{d}\int_{\mathcal{H}} p(\mathbf{h},m_i=t)D_{KL}(p(\cdot|\mathbf{h},m_i=t)||p(\cdot|\mathbf{h}))dh \\
&+ \sum_{t\in\{0,1\}}\sum_{i=1}^{d}\int_{\mathcal{H}} p(\mathbf{h},m_i=t)\log(p_m(m_i=t|\mathbf{h}))dh + C
\end{aligned}
$$

$C$ denotes a constant, therefore, the loss function is minimized if $D_{KL}(p||q) = 0$, that is, $p(x|\mathbf{h}, m_i = t) = p(x|\mathbf{h})$
